# Hemodynamic evaluation of chemical mediators of sepsis during systemic inflammatory response in an experimental animal model

**DOI:** 10.1101/2021.09.24.461714

**Authors:** Guilherme de Souza Vieira, Fernanda Antunes, Josias Alves Machado, Isabella Cristina Morales, Priscilla Olivieri Benck de Jesus, Alexandra de Faria do Amaral, André Lacerda de Abreu Oliveira

**Affiliations:** Laboratory of Physisiology and Experimental Pharmacology of the Unit of Animal Experimental, Universidade Estadual do Norte Fluminense Darcy Ribeiro, Campos dos Goytacazes, Rio de Janeiro, Brazil; Laboratory of Complexs Surgerys of the Unit of Animal Experimental, Universidade Estadual do Norte Fluminense Darcy Ribeiro, Campos dos Goytacazes, Rio de Janeiro, Brazil

## Abstract

The early diagnosis of sepsis increases the chances of its successful treatment. Biomarkers are able to distinguish between systemic inflammatory response syndrome and sepsis and are used to monitor pro- and anti-inflammatory changes associated with the host response to pathogens. A total of 11 rats underwent sepsis induction and measured systolic, diastolic and mean arterial blood pressure. Leukocyte counts, procalcitonin, and nitric oxide also were measured 0, 2, and 4 hours after the induction of sepsis using the cecal ligation and puncture method. The animals were divided into two groups: control (SHAM) and induced. Procalcitonin levels remained within the normal range for an inflammatory response throughout the experiment. There was a statistically insignificant increase in nitric oxide levels. All animals showed increased diastolic arterial blood pressure; however, the increase in the induced animals was even more pronounced. Procalcitonin and nitric oxide levels can increase due to surgical manipulation, while arterial blood pressure was not a good predictor for the onset of sepsis during the time period studied here.

## Introduction

Sepsis is a syndrome caused by an uncontrolled systemic inflammatory response of the individual that is of bacterial, fungal, or viral origin. If not treated promptly, sepsis progresses to septic shock, which is characterized by severe depletion of intravascular volume and cellular hypoxia and can lead to multiple organ failure and death [1].

With a mortality rate of 30–50%, severe sepsis and septic shock are the major causes of admission and death in the intensive care unit (ICU). In the United States, 751,000 cases and 215,000 deaths annually are estimated. In Brazil, the incidence is 57 per 1000 patients per day and the mortality rates of patients with severe sepsis and septic shock are 47.3% and 52.2%, respectively [2]. Although the symptoms are known, distinguishing the cause of sepsis in patients with clinical signs of acute inflammation in the emergency room and ICU remains problematic [3].

The early recognition and treatment of sepsis contributes to recovery success and, consequently, higher survival rates [1]. The presence of certain components in the pathogen membrane induces the release of specific inflammatory mediators characterizing the initial phase of sepsis. Biomarkers are able to distinguish between SIRS and sepsis, and a strategy to monitor the pro- and anti-inflammatory changes associated with the host response to pathogens [3].

In recent years, researchers have consequently attempted to diagnose sepsis early and change or interrupt its course. However, the poor clinical outcome and/or continuing high mortality rates of patients with sepsis have not yet resulted in an immediate or successful solution to this problem.

The use of biomarkers has been suggested to assist with early diagnosis; therefore, it may be an immediate appropriate therapy for patients with sepsis in the ICU. A biomarker is an indicator of normal biological processes as well as the pathogenic or pharmacological responses that may lead to a therapeutic intervention [4].

Procalcitonin (PCT) is a biomarker encoded by the *CALC-1* gene located on chromosome 11. The mRNA is translated into pre-PCT and the product of this translation is modified in PCT, a 116-amino-acid prohormone that is then converted into calcitonin, an active 32-amino-acid hormone that is involved in calcium and phosphorous metabolism [5].

In healthy patients, PCT is secreted almost exclusively by thyroid C-cells [5]; under these conditions, the serum PCT concentrations are very low (0.05 ng/mL) [6-7]. In the case of septicemia, particularly when associated with bacteremia, an alternative pathway for PCT production may become activated that increases its serum levels. Lung, colon, and spleen tissues can produce PCT in these cases [5]. Levels > 2 ng/mL indicate susceptibility to developing severe sepsis or septic shock [7].

PCT is considered a good clinical marker due to its high specificity and sensitivity (8), and its serum levels increase within 3 hours, peaking at around 6–12 hours in addition to being highly stable in the serum and plasma [7]. It also has a half-life of 24–30 hours in the circulation [9].

Endogenous nitric oxide (NO) is generated from L-arginine by the action of the enzyme NO synthase (NOS). The activity of NOS in the oxidation of L-arginine leads to the production of NO and L-citrulline [10].

NOS enzymes are essential to NO production, and three isoforms have been described: constitutive NOS (cNOS), which can be endothelial (eNOS) or neuronal (nNOS), both of which are calcium-dependent and inducible NOS (iNOS), which is calcium-independent [11].

Under normal physiological conditions, cNOS is present in endothelial cells, brain cells, and platelets and synthesizes NO from L-arginine in a calcium- and NADPH-dependent manner.

NO acts as a neurotransmitter when produced by nNOS expressed in the central and peripheral nervous systems, kidneys, skeletal muscles, myocardium, and pancreas [11]. It has an important physiological function as a neurotransmitter as well as a mediator in coupling neural metabolism and cerebral blood flow [10].

Not expressed under normal conditions, iNOS is induced by cytokines and/or endotoxins in several cell types including macrophages, neutrophils, Kupffer cells, and hepatocytes [12]. Under these conditions, NO is produced from L-arginine by the action of iNOS [10]. This isoform requires several hours to be expressed, but once synthesized, it releases larger amounts of NO than eNOS and nNOS and its production continues indefinitely until L-arginine and/or the co-factors required for its synthesis are depleted or when cell death occurs [12].

The expression of iNOS results in a localized or diffuse inflammatory response results from infection or tissue damage. High concentrations of NO, which is toxic to microbes, parasites, and tumor cells, can also harm surrounding healthy cells since this mechanism is responsible for the majority of inflammatory and autoimmune processes [12].

The first biological function attributed to NO was vasodilation mediation, although today NO is also believed to have a role in body temperature control during physiological and pathological conditions [13].

The wistar rat is widely used in animal experimental work because it is already known the anatomy, physiology and behavior of these animals. In addition, this animal is easy to handle and their physiological and genetic characteristics are similar to humans [14].

The objective of this study was to evaluate arterial blood pressure and the biomarkers in an experimental model of sepsis. Our hypothesis was that arterial blood pressure could predict the onset of sepsis via correlation of its value with the biomarkers involved to acquire a quick and early diagnosis.

## Material and Methods

The tests were performed on 11 male wistar rats (*Rattus norvegicus*) from the Unit of Animal Experimental da Universidade Estadual do Norte Fluminense Darcy Ribeiro, weighing between 250 and 300 grams. They were kept in appropriate cages in groups of 5 animals, covered with wood shavings. The temperature controlled environment possessed 19 °C and humidity of 50 to 60%, maintained in light/dark cycles of 12 hours. Food and water were provided *ad libitum*. At the moment of the experiment, these animals were placed alone in similar boxes, but smaller, to evaluation in the recommended times.

Anesthesia was delivered with the help of an inhalant mask made from a PET bottle and held together with a smaller mask produced from a 20 mL syringe and filled with native lime to minimize the CO_2_ rebreathing. The anesthesia was maintained with halothane throughout the procedure.

Invasive arterial blood pressure was measured by cannulation of the carotid artery in which a heparinized cannula was positioned and fixed in the carotid and fixed on the skin of the dorsal region of the neck for subsequent evaluations. The cannula was connected to the sensor of the BioAmp equipment (ADInstruments, São Paulo, Brazil) that transforms the blood pressure information and amplifies the signal in the form of graphics for the computer to enable posterior data analysis using LabChart 7 software (ADInstruments).

The animals were randomly divided into control (SHAM) and experimental groups.

In the SHAM group (n = 5), a laparotomy was performed and the cecum was exposed and returned to the abdominal cavity without causing sepsis. In the experimental group (n = 6), sepsis was induced in the animals by laparotomy and exposure of the cecum followed by cecal ligation and puncture (CLP). The cecum was returned to the abdominal cavity, which was closed with non-absorbable nylon 3/0 wire and simple interrupted sutures.

The CLP was performed as described by Witcherman et al. [15] with a slight modification. The rats were anesthetized with halothane, and a midline abdominal incision was made to expose the cecum, followed by a loose ligation of the apex of the cecum that was filled with feces. The cecum was then punctured three times with a hypodermic needle 40 × 16 and returned to the abdominal cavity to promote fecal extravasation and result in CLP-induced peritonitis.

The animals were evaluated at three time points: immediately after, 2 hours after, and 4 hours after the induction of sepsis.

The evaluation consisted of invasive blood pressure measurement, collection of 1 mL of blood for leukocyte count (WBC) and quantification of PCT and NO using a Rat Procalcitonin (PCT) Elisa Kit and a Nitric Oxide (NO_2_/NO_3_) Detection Kit. After each blood collection, the volume was replaced with 0.9% saline solution to minimize the possible effects of hypovolemia. At the end of the 4-hour evaluation, the animals were sacrificed.

The data were compiled and statistical significance was established using one-way analysis of variance (ANOVA) followed by the Newman-Keuls test (p < 0.05). The data were analyzed using GraphPad Prism^®^ 5.0.

### Ethics Statement

This study was submitted to Use Ethics Committee of animals of the Universidade Estadual do Norte Fluminense by the number 161.

## Results

The recovery of the anesthetic animals was normal and the tests could be performed at the indicated time points.

In both groups, the animals’ WBC counts did not differ significantly, but a pattern of leukocytosis at time point 2 and leukopenia at time point 3 could be observed (Fig. 1).

**Fig. 1.**
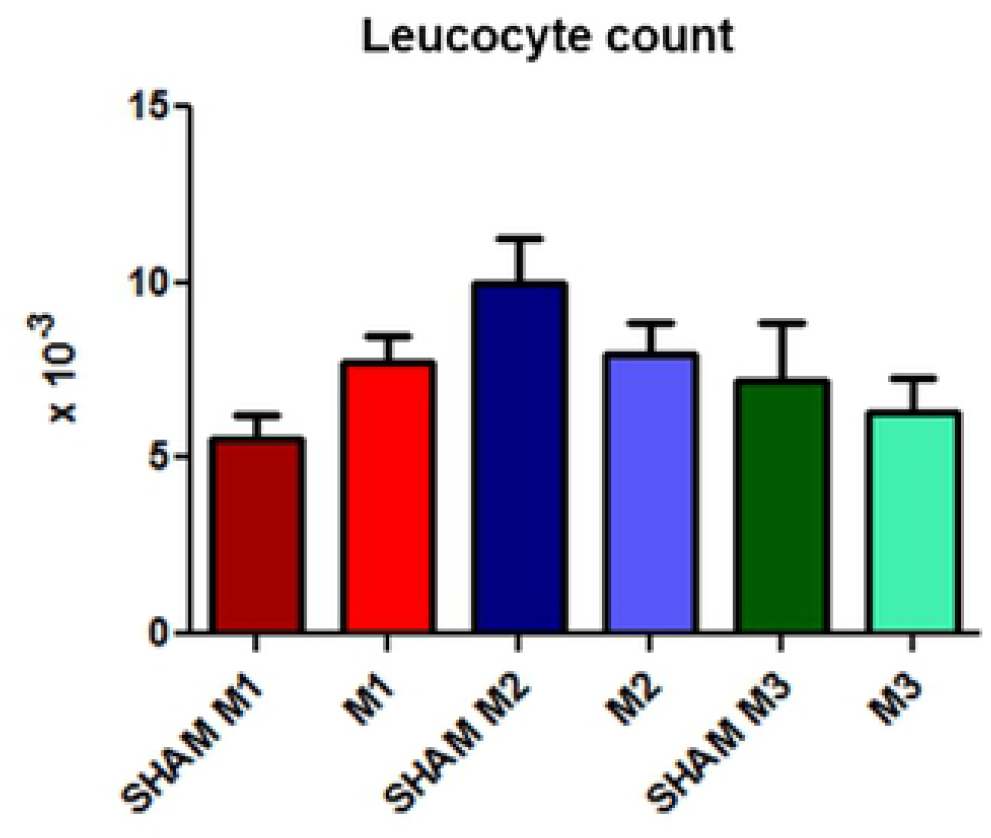
Leukocyte counts at the three time evaluated points (M1, immediately after; M2, 2 hours after; and M3, 4 hours after the induction of sepsis) compared with those in the SHAM group.

The PCT values remained very similar between groups with values close to 2 ng/mL (Fig. 2).

**Fig. 2.**
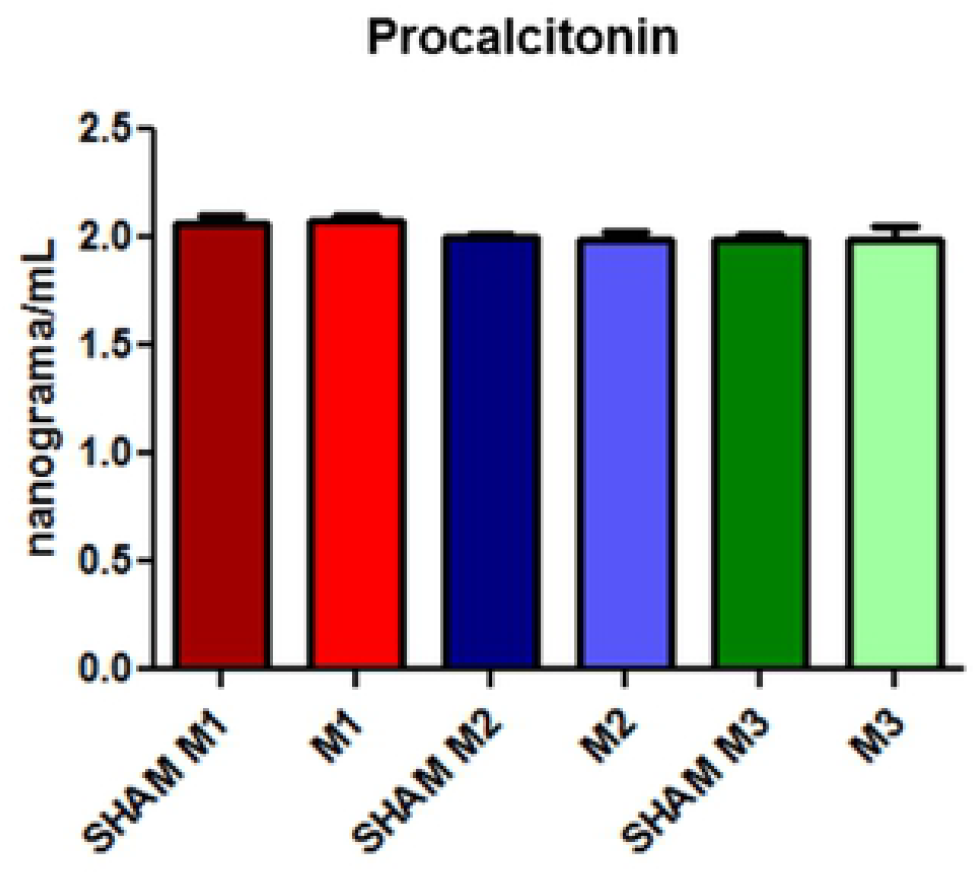
Procalcitonin concentrations at the three evaluated time points (M1, immediately after; M2, 2 hours after; and M3, 4 hours after the induction of sepsis) compared with those in the SHAM group.

The NO levels showed progressive but insignificant increases at each time point in both groups; in the induced group, the increase was even more pronounced (Fig. 3).

**Fig. 3.**
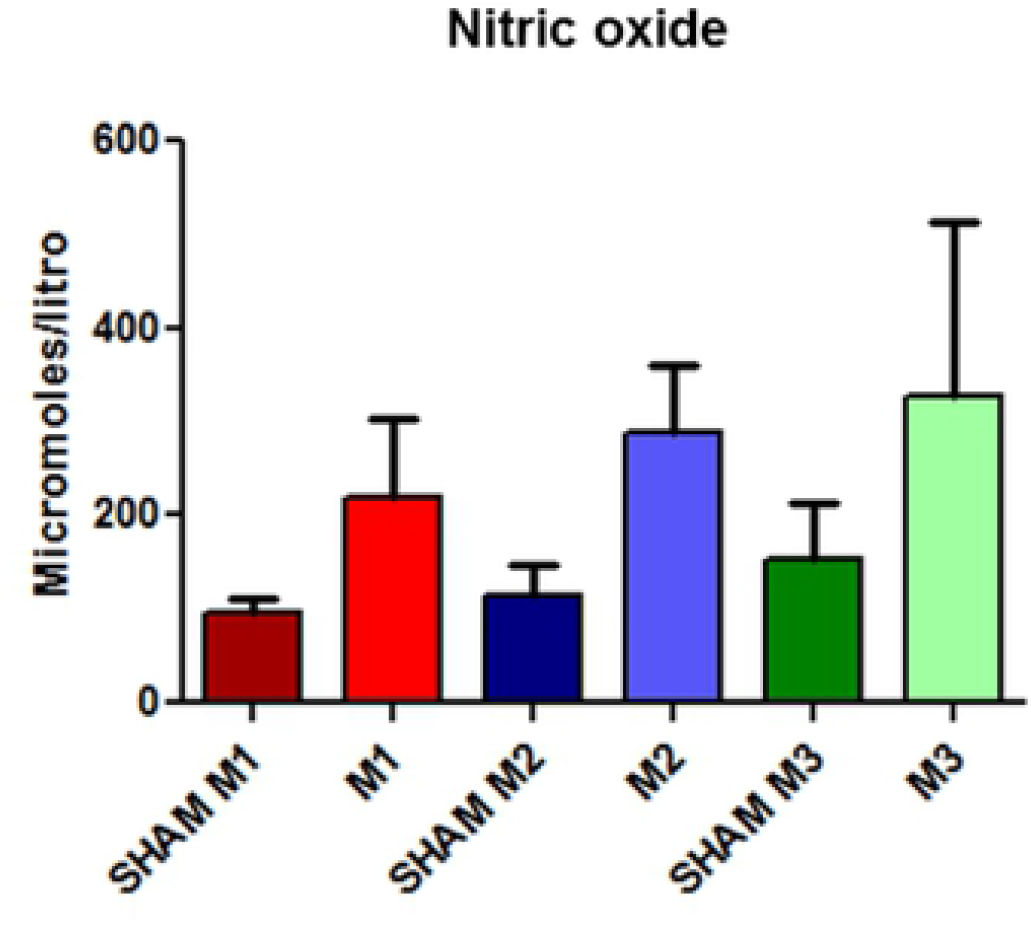
Released nitric oxide concentrations in SHAM animals and in the experimental model of induced sepsis at the three evaluated time points (M1, immediately after; M2, 2 hours after; and M3, 4 hours after the induction of sepsis).

**Fig. 4.**
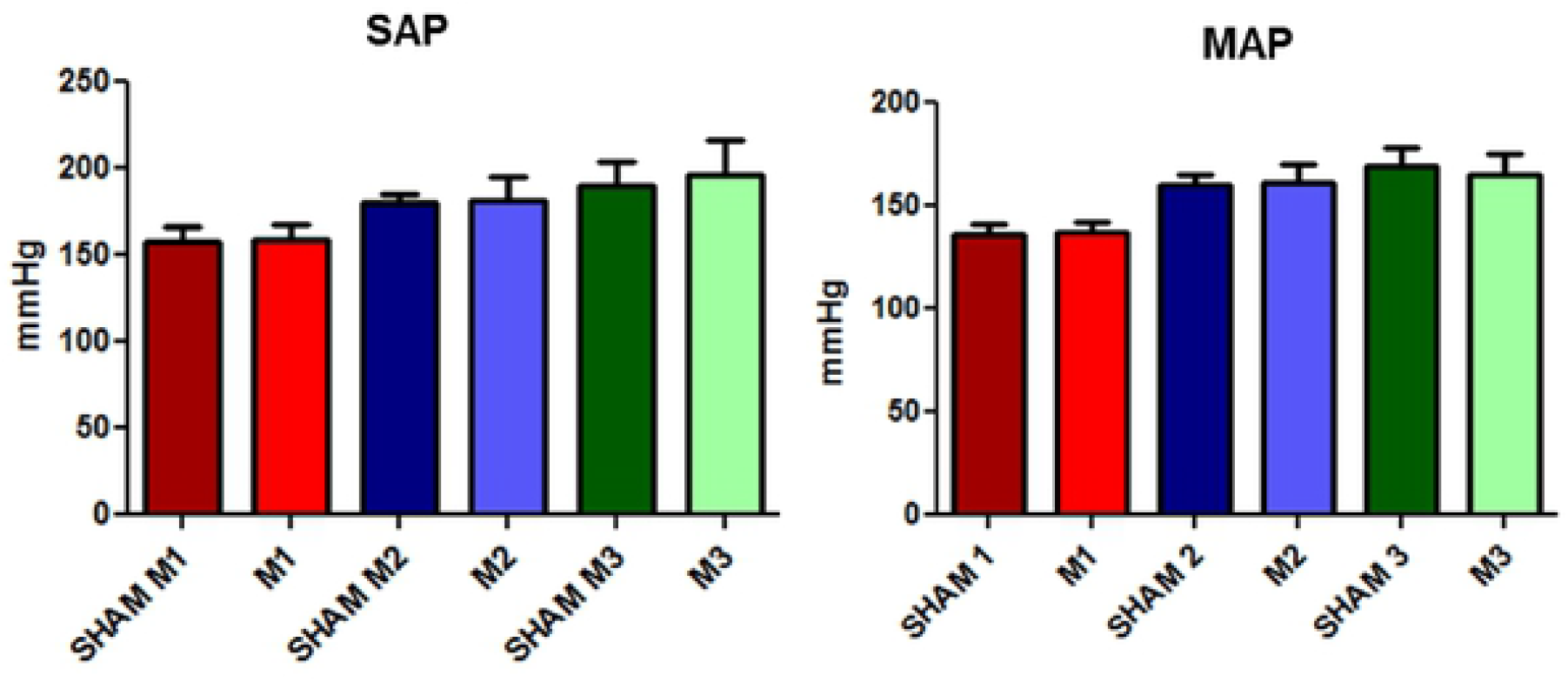
Changes in the systolic and mean arterial blood pressure in the SHAM and induced animals at the three evaluated time points (M1, immediately after; M2, 2 hours after; and M3, 4 hours after the induction of sepsis).

**Fig. 5.**
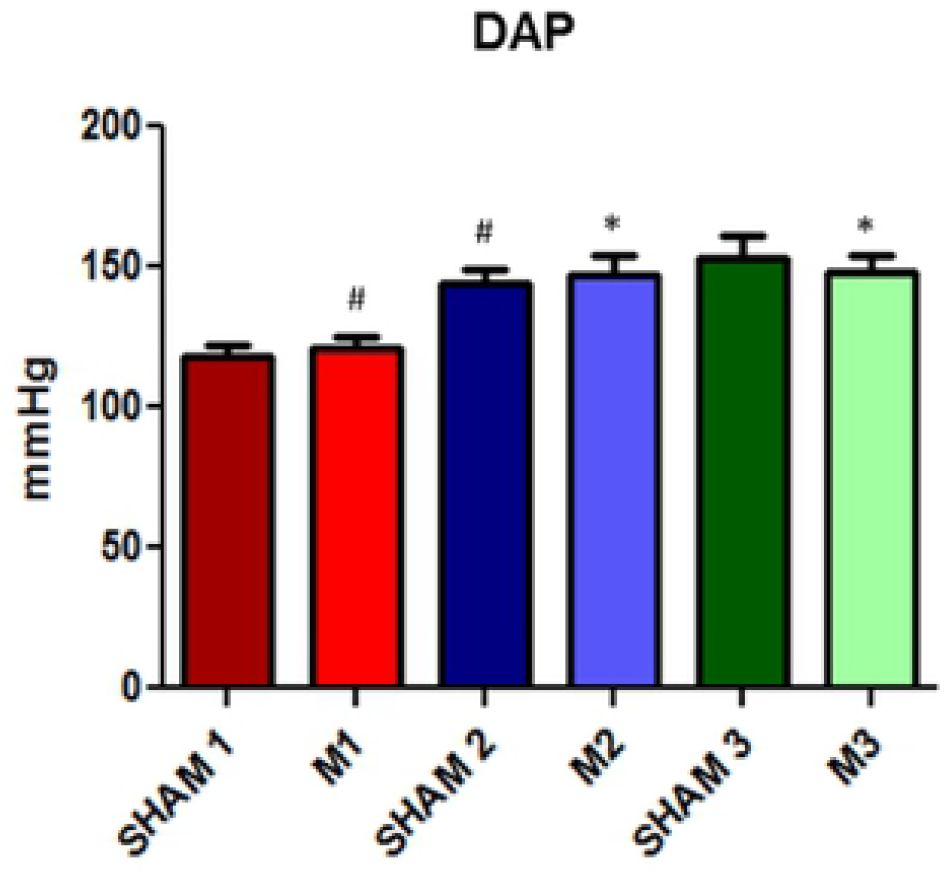
Evolution of diastolic arterial blood pressure in SHAM and induced sepsis animals at the three evaluated time points (M1, immediately after; M2, 2 hours after; and M3, 4 hours after the induction of sepsis).

Systolic arterial blood pressure (SAP) and mean arterial blood pressure (MAP) values did not differ significantly between groups. However, when comparing diastolic arterial blood pressure (DAP), a statistically significant increase in DAP was observed between the animals of the control group at time points 1 and 2 in addition to the difference between sepsis-induced animals at time points 2 and 3 compared to time point 1.

## Discussion

The PCT levels at the different time points in the induced animals did not differ significantly from the levels of the respective SHAM animals. In this case, the animals were not considered septic since the values obtained were below or very close to 2 ng/mL. However, the animals were in SIRS at all time points as identified by Arkader [6] and Liu et al. [7], who defined PCT values of 0.05–2 ng/mL as indicative of this syndrome.

The SHAM animals that underwent only cecum manipulation without CLP also had elevated PCT levels indicative of SIRS. Since these animals underwent the cannulation procedure to record the blood pressure, it is possible that these increased levels of PCT are the result of surgical manipulation as described by Diaz [16] and Soreng et al. [5], who claimed that surgeries, severe trauma, and burns are also capable of increasing PCT levels.

The animals were evaluated for only 4 hours, so it is possible that the PCT values were not yet at its peak plasma concentration. According to Liu et al. [7], PCT levels in humans increase over a period of 3 hours and peak at 6–12 hours. The time frame for this response in rats was estimated to be 4 hours since this species has a higher metabolic rate than humans. Accordingly, we estimated that the selected time frame would be sufficient for determining sepsis.

The NO levels did not show statistically significant differences between groups, although it is possible to observe an increase in both SHAM and induced animals since NO values were even higher in the last group. This demonstrates that surgical manipulation may also increase the NO plasma levels and that this model for inducing sepsis was able to increase those levels compared to the SHAM group at the three evaluated time points. The increase of NO confirms the data described by Vieira [12], who affirmed that NO levels increase the production of iNOS when there is an inflammatory involvement resulting from infection or tissue damage.

Although there were no statistically significant differences in the SAP and MAP between the SHAM and sepsis-induced animals, it was clinically possible to observe an increase in SAP, MAP, and DAP at time point 2. This difference may be due to the fact that time point 1 was immediately after the surgical procedure, when the animals could still be recovering from the anesthesia.

The maintenance of a stable MBP throughout the entire procedure shows that despite the connection between NO levels and vasodilation in septic patients, its increase in circulation did not correlate to the decrease in MAP. It can be inferred that an increase in NO precedes clinical hypotension in patients. Pereira et al. [13] also did not observe a relation between mean hypotension and NO increase in sepsis, severe sepsis, or septic shock in dogs.

DAP showed an increase between SHAM time points 1 and 2, induced time points 1 and 2, and induced time points 1 and 3. According to Hajjar et al. [17] the diastolic dysfunction induced by sepsis is not yet well established but occurs by itself or in association with systolic dysfunction. The increase in diastolic pressure observed in this study may be caused by the restriction imposed by the pericardium, which tends to promote rapid ventricular filling [18].

Moreover, Bouhemad et al. [19] evaluated the hearts of patients with septic shock using Doppler echocardiography and observed a reduced relaxation of the left ventricle in 20% of 54 patients.

Rapid ventricular filling and reduced left ventricular relaxation may have been decisive factors for the observed increase of DAP. Since cardiac evaluation, troponin level measurements, and echocardiography were not performed in this case, it is not possible to confirm that this finding may be explained by the changes mentioned above.

Munt et al. [20] also described that more pronounced diastolic changes were observed in the group of septic patients that did not survive.

## Conclusions

After the end of the experiment and data analysis, we can conclude that despite PCT being a recommended biomarker for the diagnosis and prognosis of sepsis, its production may be increased for other causes such as surgeries and trauma rather than bacterial infections.

The plasma NO concentration may be increased due to surgical manipulation. Despite the fact that this biomarker promotes vasodilation, it is not related to mean arterial hypotension.

The use of arterial blood pressure as the only method of assessment was not a good predictor of sepsis onset during the 4-hour study period. The results of diastolic hypertension found in the present study are not consistent with those found in the studies that cite hypotension in cases of septic shock.

